# Biliverdin Reductase A is a major determinant of neuroprotective Nrf2 signaling

**DOI:** 10.1101/2025.06.04.657936

**Authors:** Chirag Vasavda, Ruchita Kothari, Navneet Ammal Kaidery, Suwarna Chakraborty, Sunil Jamuna Tripathi, Ryan S. Dhindsa, Samaneh Saberi, Julia E. Lefler, Priyanka Kothari, Kalyani Chaubey, Cristina Ricco, Adele M. Snowman, Michael C. Ostrowski, Eugenio Barone, Lakshminarayan M. Iyer, L. Aravind, Sudarshana M. Sharma, Andrew A. Pieper, Bobby Thomas, Solomon H. Snyder, Bindu D. Paul

## Abstract

Biliverdin reductase A (BVRA), the terminal enzyme in heme catabolism, generates the neuroprotective and lipophilic antioxidant bilirubin. Here, we identify a novel non-enzymatic role for BVRA in redox regulation. We show that BVRA directly interacts with nuclear factor erythroid-derived factor-like 2 (Nrf2), the master regulator of redox homeostasis, to modulate target signaling pathways. ChIP-seq and RNA-seq analyses reveal that this interaction coordinates the expression of neuroprotective genes that are typically dysregulated in Alzheimer’s disease and other neurodegenerative conditions. Thus, this previously unknown BVRA-Nrf2 axis controls an essential pathway of redox signaling in neuroprotection. Our findings establish BVRA as a dual-function integrator of antioxidant defenses in both the lipophilic and hydrophilic subcellular compartments, bridging these two distinct and critical cellular protection mechanisms in the brain. This advancement in understanding the endogenous antioxidant system of the brain positions the BVRA-Nrf2 axis as a promising therapeutic target for neurodegenerative disease.

**Significance Statement:** We show a non-canonical role for biliverdin reductase A (BVRA), classically known as the biosynthetic enzyme for bilirubin, in nonenzymatic modulation of antioxidant neuroprotective nuclear factor erythroid-derived factor-like 2 (Nrf2) signaling in the brain. Both BVRA and Nrf2 signaling are compromised in neurodegenerative diseases such as Alzheimer’s disease, and the BVRA-Nrf2 axis offers a new direction for developing neuroprotective therapies.

## Introduction

Biliverdin reductase A (BVRA), encoded by the *Blvra* gene, and the terminal enzyme in heme catabolism and the major biosynthetic enzyme for bilirubin in adults (Fig. 1*A*) (1, 2). Bilirubin is a potent lipophilic antioxidant with protective efficacy in advanced cardiovascular disease associated with hyperbilirubinemia (3–9), as well as in malaria (10) and metabolic disorders (11, 12). Beyond its canonical role, BVRA exhibits additional diverse enzymatic and non-enzymatic functions, modulating key signaling pathways such as focal adhesion kinase signaling in the brain, insulin signaling, phosphoinositide 3-kinase (PI3K)/Protein kinase B (Akt) signaling, and mitogen-activated protein kinase (MAPK) cascades (11, 13–21). Interestingly, at near-neutral pH BVRA utilizes NADH as a cofactor, whereas it uses NADPH at alkaline pH, enabling flexibility of function under different cellular conditions (22–24). Importantly, while BVRA’s roles in peripheral tissues are well-documented, its functions in the central nervous system are not as well understood, despite its significant expression in brain (2).

**Fig. 1.**
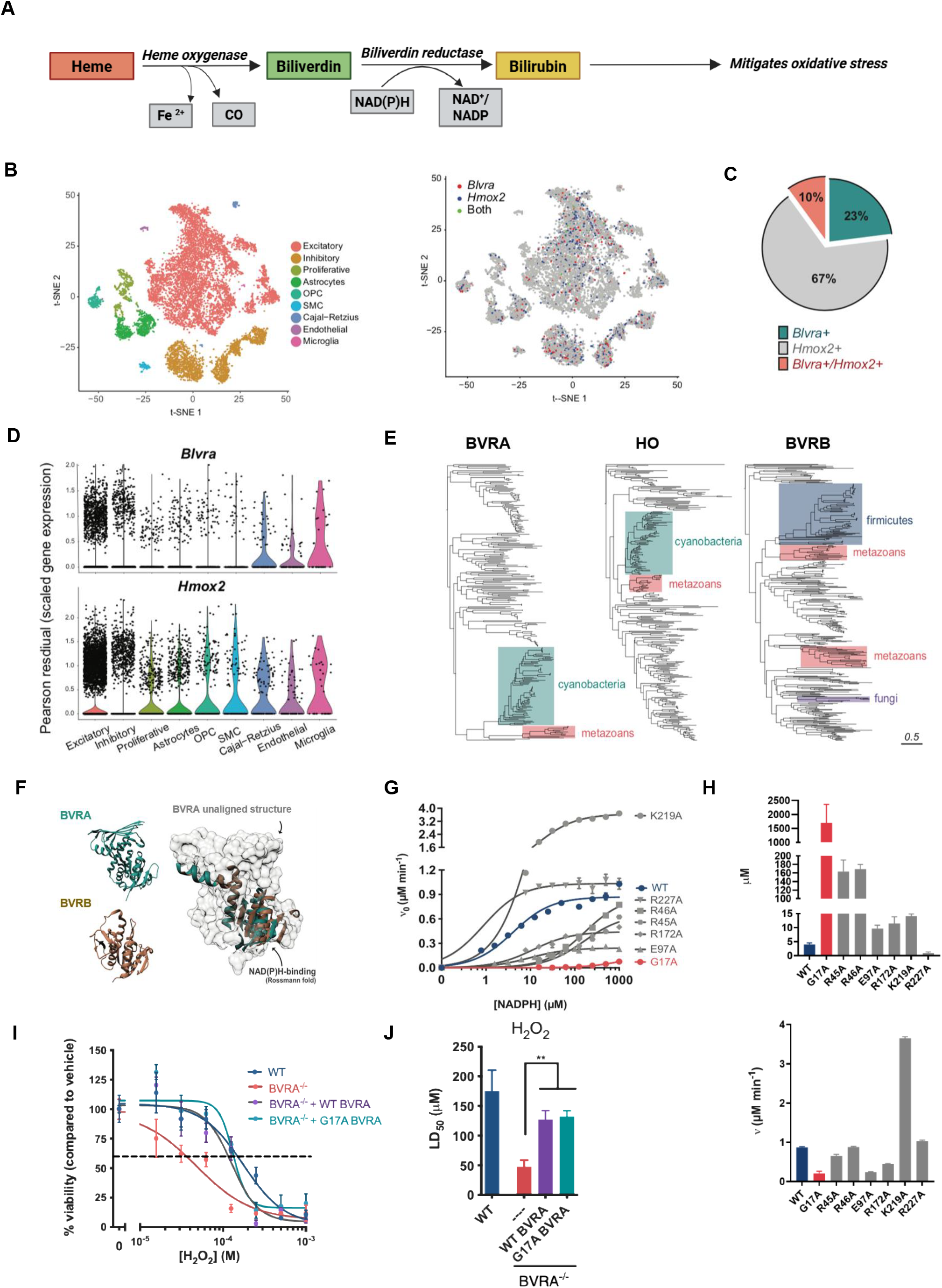
BVRA exhibits expression profiles and redox homeostatic activity unlinked to heme catabolism. (*A*) Schematic representation of the heme catabolic pathway. Heme derived from hemoglobin from senescent red blood cells is metabolized by heme oxygenases (HO) to produce the green pigment, biliverdin. Biliverdin reductase A (BVRA) then reduces biliverdin to produce the yellow pigment, bilirubin, which is a potent antioxidant. (*B*) T-distributed stochastic neighbor embedding (t-SNE) analysis of single-cell RNA expression in primary neuronal cultures from WT C57BL/6J mice. Each point represents a single cell. Left panel: Cells are clustered by gene expression into populations with similar cell type and function. Right panel: Expression distribution of *Blvra, Hmox2*, both individually and together, across cell population clusters. (*C*) Quantified percentage of total cells expressing *Blvra, Hmox2*, and a combination of both genes. (*D*) Scaled Pearson residuals of *Blvra* and *Hmox2* gene expression in different cell populations. A residual >1 indicates a higher observed to expected frequency ratio. Each point represents a single cell. (*E*) Phylogenetic trees of BVRA, HO, and BVRB respectively. (*F*) Protein structures of BVRA and BVRB obtained from Kavanagh et. al. (PDB Entry - 2H63 (pdb_00002h63) and Periera et. al., respectively (54). Structural comparison of the two BVR proteins aligned at the Rossmann fold of the NADPH-binding domain. (*G*) Activity of WT BVRA and BVRA mutants measured at varying concentrations of NADPH and 10 μM biliverdin. Activity measured by absorbance reading of bilirubin production at 442 nM with V_o_ calculated according to the method by Michaelis and Menten. n = 3 in triplicate. (*H*) Quantified V_max_ and K_m_ of WT BVRA and BVRA mutants. (*I*) Viability of WT and BVRA MEFs transfected with empty vector, WT BVRA, and G17A-BVRA after 8 h of exposure to serial dosing of H_2_O_2_. Data is normalized to viability of vehicle condition. (*J*) Quantified LD_50_s (I) of H_2_O_2_ exposure in WT and BVRA MEFS. n = 3 in triplicate.

We previously showed that both bilirubin and BVRA are highly abundant in the lipid-rich brain (2). Bilirubin, being lipophilic, functions in a manner complementary to glutathione (GSH), the abundant hydrophilic antioxidant to confer beneficial effects (2, 7–9, 25). Bilirubin more effectively prevents lipid peroxidation than soluble cellular components, such as protecting cytosolic proteins from oxidation (2, 8). Bilirubin is also a potent scavenger of superoxide (O_2_^•−^) radicals, with mice lacking BVRA displaying elevated susceptibility to oxidative stress (2, 9). We have shown that bilirubin is a potent scavenger of O_2_^•−^ and cannot neutralize other forms of reactive oxygen species (ROS) such as H_2_O_2_ and the hydroxyl radical (OH^•^)(2).

Despite bilirubin’s apparent lack of reactivity towards these ROS, *Blvra*^−/−^ mice are still hypersensitive to H_2_O_2_ and complementing cells with BVRA is protective, suggesting that these mice may be deficient in a mechanism dependent on BVRA, but not bilirubin to counteract oxidative stress (2). Here, we reveal a redox-regulatory role of BVRA that entails a non-enzymatic interaction with the nuclear factor erythroid-derived factor-like 2 (Nrf2) transcription factor, which is well known as the master regulator of antioxidant responses (26). We demonstrate that direct binding of BVRA to Nrf2 enhances transcriptional regulation of neuroprotective gene networks implicated in Alzheimer’s disease (AD) and neurodegeneration. This BVRA-Nrf2 axis establishes a novel link between heme metabolism and physiologic redox homeostasis, revealing opportunities to therapeutically bolster endogenous neuroprotection.

## Results

### Heme-independent expression patterns and functional roles of brain BVRA

The first enzyme in heme catabolism in the brain, heme oxygenase 2 (HO-2) accounts for most of the HO activity in the nervous system and, like HO-1, regulates redox balance (27, 28). HO-2 deletion aggravates oxidative stress induced by seizures, glutamate, and inflammation, and also causes cerebral vascular injury (29). To determine whether BVRA operates independently of the heme oxygenase (HO) system, we first identified brain cell types in which heme metabolism is crucial. Using single-cell RNA sequencing (scRNA-seq), we compared *Blvra* (which encodes BVRA) and *Hmox2* (which encodes HO-2) expression profiles in whole-brain lysates. Both *Blvra* and *Hmox2* were detected in numerous cell types in the brain (Fig. 1*B*). On the other hand, *Hmox1* mRNA (which encodes HO-1) is inducible (30) and not expressed at baseline, rendering it undetectable in most cells analyzed. Notably, *Blvra* exhibited more restricted cellular expression than *Hmox2*, with HO-2 mRNA levels nearly tripling those of *Blvra* transcripts. Strikingly, 23% of brain cells expressed *Blvra* mRNA without detectable *Hmox2* mRNA and only 10% expressed both *Blvra* and *Hmox2* (Fig. 1*C*), a pattern observed broadly across cell types and brain regions (Fig. 1*D*). This surprisingly broad presence of BVRA in the absence of HO-2, is puzzling and suggests a potential function independent of heme metabolism. This dissociation, corroborated by independent datasets (Allen Brain Atlas, Protein Atlas) (31), implies that BVRA operates in contexts unrelated to bilirubin generation.

We next examined the evolutionary histories of these principal enzymes involved in heme degradation: the BVRs (BVRA, BVRB), and HO. BVRA and BVRB are distinct enzymes with BVRA being the main bilirubin producing enzyme in adults and BVRB in the fetal stages (32). BVRA belongs to the expansive glyceraldehyde 3-phosphate dehydrogenase (GAPDH)-like clade of redox enzymes, whose members feature an N-terminal NADPH/NADH-binding Rossmann fold catalytic domain and a C-terminal GADC domain responsible for substrate selection and binding. Structural studies of the related *Synechocystis* BVRA revealed that its C-terminal domain accommodates two biliverdin molecules via π-π stacking, both of which are believed to participate in the redox reaction. Based on the *Synechocystis* BVRA structure and its interactions with NADPH and biliverdin, and the sequence conservation pattern, we identified several residues for further mutational studies (G17, E97, R172, R46, R227, K219). We also noticed a cysteine residue (C292) that is conserved across vertebrates and some cyanobacteria, which may play a role in BVRA-mediated redox homeostasis (Fig. S1). In contrast, BVRB lacks the GADC domain and is comprised of only the NADPH-binding Rossmann fold catalytic domain, which is also present in BVRA (Fig. 1*F*). Consistent with the absence of a dedicated substrate-binding domain, it exhibits greater substrate promiscuity. HO, on the other hand, is a heme-dependent all–α-helical oxygenase that is structurally unrelated to BVRA and BVRB.

Given that homologs of these enzymes are reported in prokaryotes, we explored their phylogenetic affinities and genomic contexts to gain insights into the evolution of this pathway. Phylogenetic analyses show that both BVRA and HO are ultimately of cyanobacterial origin. However, their patterns of acquisition by eukaryotes differ, with the gene for BVRA appearing to have been transferred from cyanobacteria to an early metazoan (Fig. 1*E*) and HO having been acquired earlier in eukaryotic evolution, as evidenced by its presence in diverse eukaryotic lineages such as oomycetes and haptophytes. BVRB, by contrast, appears to represent a lateral transfer from a firmicute lineage again into an early metazoan. Notably, homologs related to all three enzymes were transferred on multiple independent occasions from prokaryotes to eukaryotes. For example, one such paralog of BVRA in eukaryotes, the diphenol detoxifying enzyme dihydrodiol dehydrogenase (DHDH), was independently acquired from bacteria.

Furthermore, even within metazoans, the phyletic patterns of BVRA and BVRB enzymes differ considerably: for example, BVRB is widely present in insects, where HO and BVRA are largely absent. However, HO and BVRA, display a degree of concordance in their distribution among other metazoans. These patterns support the hypothesis that the heme catabolic pathway combining the roles of BVRA and HO likely evolved near the base of the animal lineage, whereas BVRB, given its broader substrate range, including flavins and ferric ions, may have been secondarily incorporated into heme catabolism in vertebrates, perhaps as an extension of its more generalized role in redox modification of exogenous toxic small molecules. The cyanobacterial versions of HO and BVRA likely functioned originally in the metabolism of phycobilins associated with proteins of their light-harvesting systems. Their early incorporation and shared distribution in animals imply that they were co-opted for the emerging importance of heme metabolism in the context of oxygen transport in multicellular animals.

To further characterize the potentially novel function of BVRA decoupled from HO, we utilized *Blvra*^−/−^ mice lacking exon 3 (encoding the NAD(P)H-binding domain) abolishing bilirubin production (2). Despite originally creating this model to interrogate bilirubin’s antioxidant effects, our earliest observations implicated BVRA itself in redox regulation, including protection against the hydrophilic H_2_O_2_ oxidant. We therefore generated a series of mutants in the NADPH and biliverdin binding domains of BVRA, including the G17A mutant described earlier (21, 33) to assess their roles in BVRA function (Fig. S1). As previously reported, a glycine to alanine mutation of residue 17 (G17A) disrupted NADPH binding and significantly diminished BVRA-mediated bilirubin production (Fig. 1*G*). Specifically, G17A conferred a 5-fold reduction in Vmax and a 300-fold increase in Km (Fig. 1*H*) of the enzyme. Furthermore, several other mutants derived from residues that are predicted to bind NADPH/NADH or biliverdin also showed diminished activities (Fig. 1*G*).

Next, we transfected BVRA^−/−^ mouse embryonic fibroblasts (MEFs) with WT or G17A-BVRA to test whether this enzymatically dead mutant could rescue cells exposed to H_2_O_2_ despite its inability to produce bilirubin (Fig. 1*I*). G17A-BVRA rescues *Blvra*^−/−^ cells to a similar extent as WT-BVRA, hinting further at a redox function of BVRA independent of bilirubin production and heme catabolism (Fig. 1*J*).

### Depleting BVRA blunts Nrf2 signaling

To assess how BVR may exert antioxidant activity independent of its role in bilirubin production, we monitored the gene expression of several antioxidant genes via unbiased whole transcriptome sequencing of WT and BVRA^−/−^ hippocampal neurons at baseline and under oxidative stress induced by H_2_O_2_. After this toxic exposure, several genes were differentially expressed in both cell types (Fig. 2*A-C*, (*SI Data Table 1*). We focused on genes that show a fold difference of at least 2 in upregulation or downregulation between the WT and *Blvra*^−/−^ genotypes. Among the genes upregulated, the cholesterol biosynthetic pathway was most prominently represented, with upregulation in BVRA^−/−^ cells under stress BVRA^−/−^ cells. Most significant among the downregulated pathways was the nuclear factor erythroid-derived factor-like 2 (Nrf2) signaling pathway (Fig. 2*D*, right panel), with Nrf2 ranking significantly high amongst the transcription factors implicated in the differential gene expression between the WT and BVRA^−/−^ cells (Fig. 2*E*). We therefore chose to focus on Nrf2 signaling for the remainder of our study to determine the specific role of BVRA in the response to oxidative stress.

**Fig. 2.**
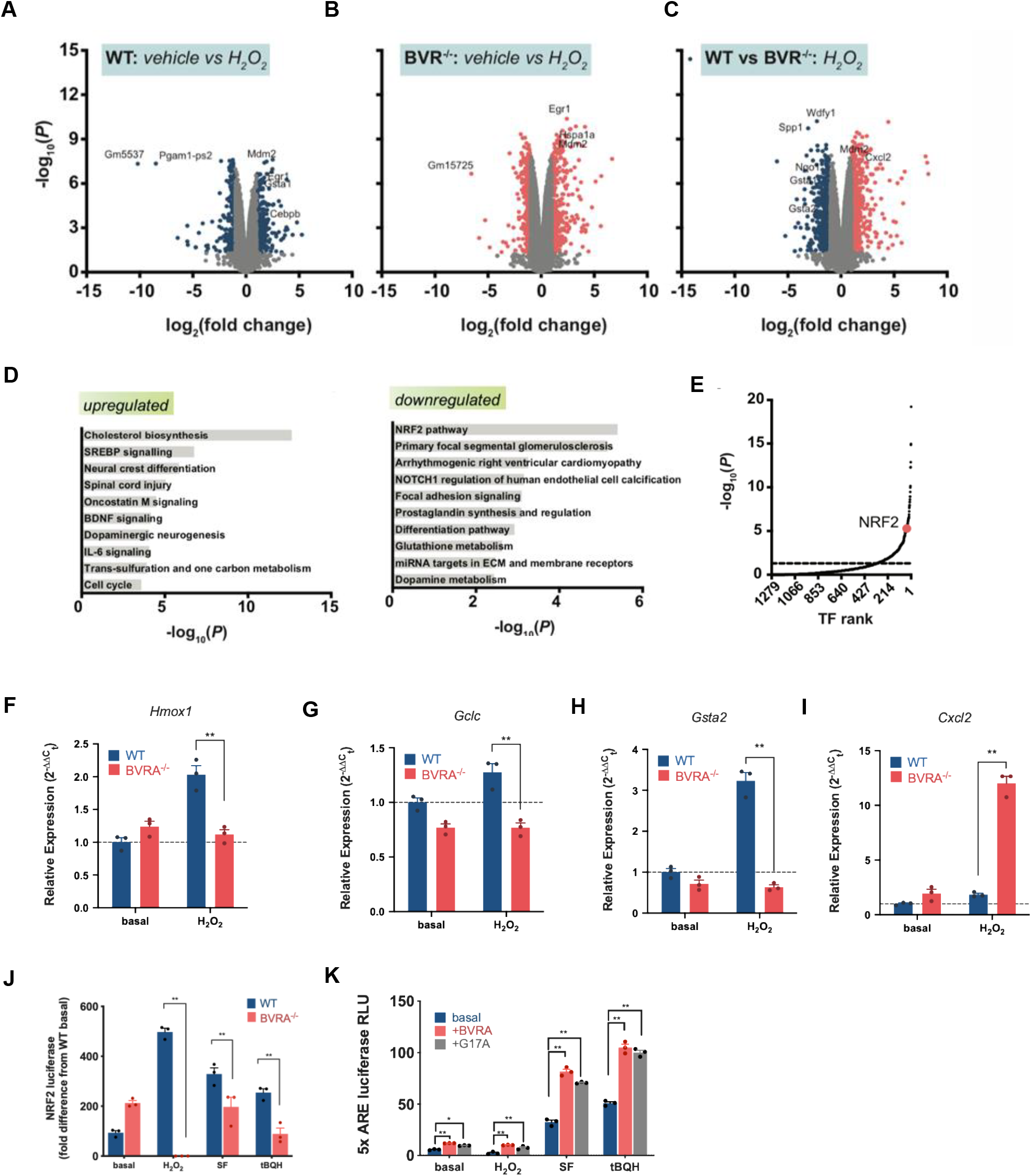
Nrf2 activity is disrupted in BVRA^−/−^ neurons and fibroblasts. (*A*) Volcano plot analysis of genes from WT and *Blvra*^*−/−*^ primary hippocampal neurons. Each plot displays the fold change of gene expression on a log scale with their corresponding P values plotted on a log scale. Each dot represents a gene of interest. Change in gene expression between WT from vehicle to H_2_O_2_ condition are shown (*B*) Change in gene expression in *Blvra*^*−/−*^ neurons from vehicle to H_2_O_2_ condition. (*C*) Change in gene expression between WT and *Blvra*^*−/−*^ neurons after H_2_O_2_ treatment. Red dots indicate genes that are upregulated with H_2_O_2_ condition in *Blvra*^*−/−*^ neurons. Blue dots indicate genes that are downregulated with H_2_O_2_ condition in *Blvra*^*−/−*^ neurons. (*D*) Logarithmically-scaled *p* values of molecular pathways that are upregulated (left panel) and downregulated (right panel) in *Blvra*^*−/−*^ neurons compared to WT controls under the H_2_O_2_ condition. Transcription factors ranked by log-scaled *p* values. A higher *p* value indicates transcription factors that are more significantly dysregulated in BVRA^*−/−*^ compared to WT neurons in the H_2_O_2_ condition. Transcription factors ranked from 1, most significant, to 1279, least significant. (*F-I*) Validation of expression of genes regulated by Nrf2 in WT and BVRA^*−/−*^ neurons. Wild-type (WT) and BVRA^*−/−*^ primary hippocampal neurons were treated with or without 200 μM H_2_O_2_ for 6 h and expression of Nrf2 targets were analyzed by qPCR. n=3, SEM, *p* <0.01, Student’s t test. (*J*) Luciferase reporter assays in WT and BVRA*−/−* MEFs transfected with Gsta4 luciferase reporter construct after exposure to vehicle, 50 μM H_2_O_2_, 10 μM SFN, and 25 μM TBHQ. n = 4 in triplicate. Sidak’s multiple comparison (*K*) Luciferase reporter assays in stable cell HEK293 cell lines harboring a 5xARE reporter constructs transfected with either WT BVRA or G17A-BVRA and after exposure to either vehicle, 200 μM H_2_O_2_, 10 μM SFN, and 25 μM TBHQ. n = 4 in triplicate. Tukey’s multiple comparison. (*L*) BVRA^−/−^ MEFs are more sensitive to erastin, an inhibitor of the XcT transporter. WT and BVRA^−/−^ MEFs were treated overnight for 18 hours with increasing concentrations of erastin, a xCT inhibitor and ferroptosis inducer, and cell viability was assessed by MTT assay. n=7, SEM, ^∗^*p < 0.05*, \*\**p <0.01*, \*\*\**p <0.001*, Student’s t test.

Nrf2 is a transcription factor that regulates the expression of several antioxidant response proteins. Under basal conditions, Nrf2 is sequestered by the Kelch-like ECH-associated protein 1 (Keap1) in the cytosol and ubiquitinated for targeted degradation by the 26S proteasome(34). In response to oxidative stress, Keap1 is modified on its reactive cysteine residues and dissociates from Nrf2, preventing its ubiquitination and proteasomal degradation (35–39). Nrf2 then translocates to the nucleus, where it binds the antioxidant response element (ARE) to orchestrate transcription of a battery of vital antioxidant genes necessary for cytoprotection. To evaluate whether Nrf2 signaling is disrupted in BVRA^−/−^ cells, we analyzed the transcript levels of Nrf2 target genes such as *Hmox1, Gclc, Gsta2* and *Cxcl2* (Fig. 2*F-I*). The expression of *Hmox1, Gclc* (which encodes the catalytic subunit of glutamate–cysteine ligase, a key enzyme in glutathione biosynthesis) and *Gsta2* (which encodes glutathione S-transferase A2, an enzyme involved in detoxification of electrophilic compounds) were blunted in the BVRA^−/−^ cells, whereas the proteins involved in the immune response and inflammation such as *Cxcl2* (encoding the chemokine (C-X-C motif) ligand 2) were upregulated.

Next, we compared the activity of the Nrf2 promoter in WT and BVRA^−/−^ cells using a luciferase reporter assay. In this dual-luciferase firefly reporter system, where the luciferase gene is regulated by the promoter for glutathione S-transferase A4 (GSTA4), a target of Nrf2. Thus, Nrf2 activity at the *Gsta4* promoter initiates luciferase transcription, which can be measured via luciferin luminescence. We quantified the activation of the *Gsta4* promoter in WT and BVRA^−/−^ MEFs in response to three Nrf2 activators: H_2_O_2_, sulforaphane (SFN), and tert-butyl hydroquinone (*t*-BHQ). SFN is a compound known to modify the reactive cysteines on Keap1, thus releasing Nrf2 before ubiquitination and increasing overall expression of intracellular Nrf2 (40). *t*-BHQ is also known to activate Nrf2, although its mechanism not well-established (41). H_2_O_2_ treatment increases luciferin reporter detection in WT cells, indicating Nrf2 activation by proxy of GST gene transcription. However, this increase is not seen in BVRA^−/−^ cells, which are unable to elicit Nrf2 activity and GST luciferase expression. The same pattern was also noted in the SFN and tBHQ treatment conditions (Fig. 2*J*).

To determine whether enzymatically dead BVRA also potentiates Nrf2 transcriptional activity, we introduced WT and G17A BVRA in a HEK293 cell line harboring an Nrf2 regulated luciferase reporter. In this HEK293 cell line, the promoter controlling luciferase expression is directly downstream of the antioxidant response element promoter that Nrf2 binds, allowing us to use luciferase expression as a readout for Nrf2 transcriptional activity in general and not limited to GSTA4 expression as in the previous experiment. WT- and G17A-BVRA both increase luciferase activity significantly and to the same degree in each treatment condition (Fig. 2*K*). Enzymatically dead BVRA was equally efficacious as WT BVRA in activating Nrf2 *in vitro*, underscoring BVRA’s novel non-catalytic Nrf2-inducing role. We next analyzed the effect of erastin, an inhibitor of the cystine transporter, one of whose subunits is encoded by *Slc7a11*, a target of Nrf2, on the viability of these cells. BVRA^−/−^ MEFs were more susceptible to increasing concentrations of erastin as compared to WT cells, further confirming that depletion of BVRA compromises Nrf2 signaling (Fig. 2*L*).

### BVRA physically and genetically interacts with Nrf2

To characterize the relationship between BVRA and Nrf2, we aimed to breed BVRA and Nrf2 double-knockout mice. In our attempts to create such mice, no *Blvra*^−/−^/ *Nfe2l2*^−/−^ pups were ever obtained, indicating that this interaction is crucial for survival and function. In the crosses conducted, we expected roughly 9% of the pups to present with the double-null genotype but were unable to obtain double knockout pups (Fig. 3*A, B*). All other possible genotypes were generated at near their expected frequencies. This finding suggests that BVRA and Nrf2 are genetically linked and that deleting both genes is embryonic lethal, confirming a genetic interaction between *Blvra* and *Nfe2l2* that is crucial for survival.

**Fig. 3.**
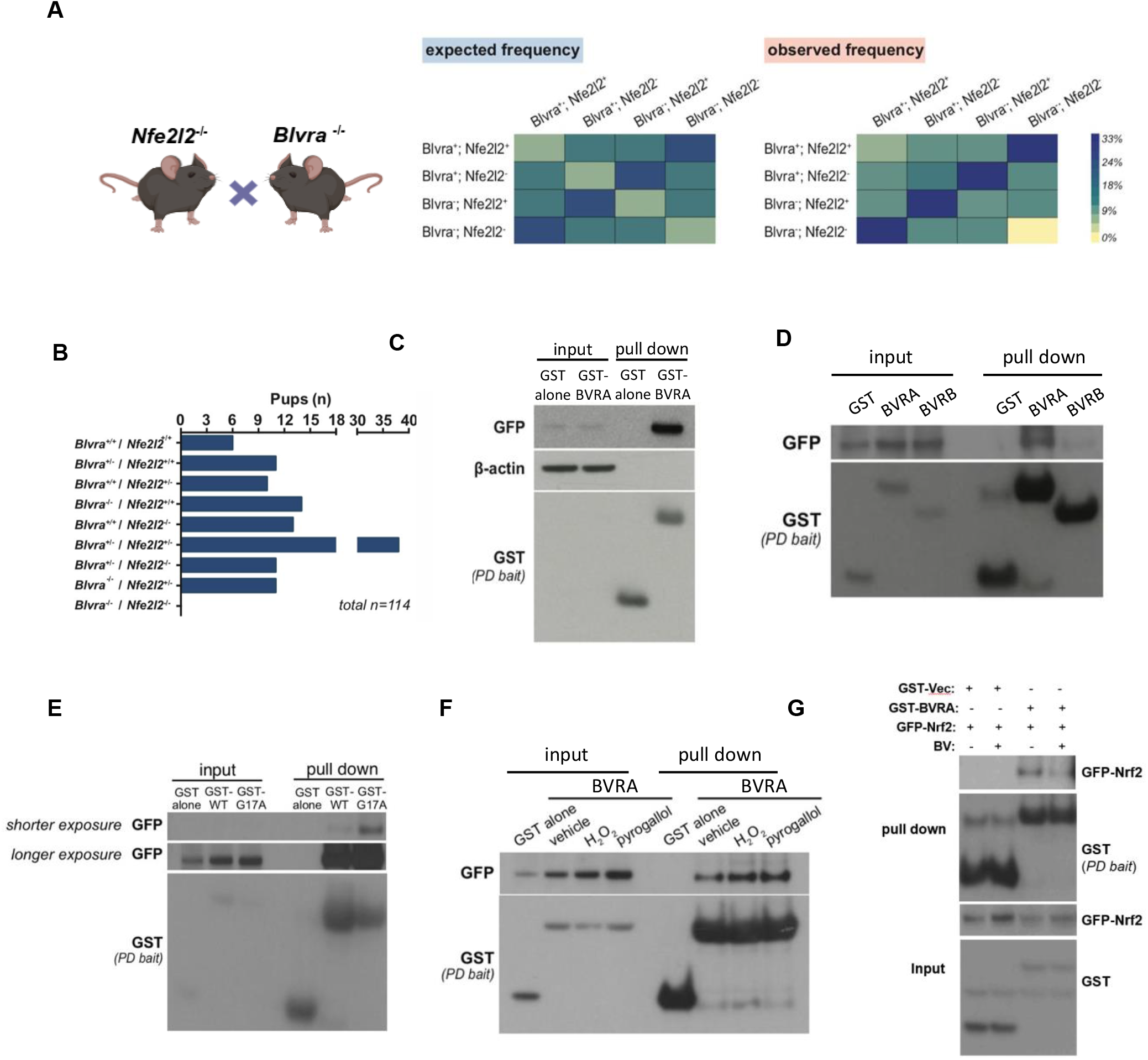
BVRA genetically and physically interacts with Nrf2. Expected versus observed frequency of pups from the *Blvra*^*−/−*^ and *Nfe2l2*^*−/−*^ (*Nrf2*^*−/−*^) cross. (*B*) Number of observed pups of each genotype. n = 114 pups total. (*C*) GST pulldown with glutathione sepharose beads of GST-BVRA and GFP-Nrf2 transfected HEK293 cells. GST-only vector control was used as a negative control. Blots were probed for GFP-Nrf2 and GST-BVRA with their respective tags. (*D*) Co-immunoprecipitation of GFP-Nrf2 with GST-BVRA and GST-BVRB. Inputs at the right indicate samples taken before incubation in beads to monitor whole cell expression of transfected proteins. Blots were probed for GFP-NRF2 and GST-BVRA with their respective tags. (*E*) GFP-Nrf2 co-immunoprecipitation with GST-WT BVRA and GST-G17A-BVRA. Shorter exposure was used for G17A IP lanes and longer exposure was used for the remaining lanes. Blots were probed for GFP-Nrf2 and GST-BVR with their respective tags. (*F*) Co-immunoprecipitation of GST-BVRA and GFP-Nrf2 with varying oxidative stress conditions, including vehicle, 200 μM H_2_O_2_, and 100 μM pyrogallol. Blots were probed for GFP-Nrf2 and GST-BVRA with their respective tags. (*G*) Interaction of GST-BVRA and GFP-Nrf2 in the presence of 10 μM biliverdin in HEK293 cells. Blots were probed for GFP-Nrf2 and GST-BVRA with their respective tags.

To determine whether BVRA interacts with Nrf2, we conducted a GST pull down assay in HEK293 transfected with constructs harboring GFP-tagged Nrf2 and either GST vector alone or a construct encoding GST-BVRA. GST-BVRA robustly bound Nrf2 unlike GST alone, as revealed by western blot analysis (Fig. 3*C*). Given that BVRA and BVRB are the two enzymes involved in bilirubin production, but have completely different evolutionary histories and are only remotely related, we analyzed whether BVRB has an Nrf2-interacting role. We simultaneously co-expressing GFP-Nrf2 with either GST-BVRB, GST-BVRA or GST alone in HEK293 cells and conducted GST pulldown assays. We observed that BVRB does not associate with Nrf2 and this function of BVR remains specific to BVRA (Fig. 3*D*). To determine whether this interaction is dependent on the BVRA’s bilirubin-producing activity, we transfected cells with Nrf2 and enzymatically dead G17A BVRA, which cannot bind the cofactors NADPH or NADH. G17A BVRA not only pulls down Nrf2 but does so more robustly than WT BVRA, indicating that this interaction is independent of its catalytic function (Fig. 3*E*).

As both BVRA and Nrf2 modulate redox homeostasis, we also analyzed whether the formation of the BVRA-Nrf2 complex increases in response to oxidative stress. We therefore simulated pro-oxidant conditions to monitor changes in the BVR-Nrf2 interaction. HEK293 cells exposed to H_2_O_2_ and pyrogallol, a superoxide generator (42), exhibit increased Nrf2 expression compared to vehicle conditions, which was associated with increased BVRA-Nrf2 complex formation via co-immunoprecipitation. (Fig. 3*F*). Notably, there was no affinity difference in interaction under different redox conditions.

We also analyzed whether BVRA influences stability of Nrf2. As Nrf2 stability and levels are regulated by ubiquitination, we first examined whether BVRA increases Nrf2 expression by binding and hindering ubiquitin association. Through co-immunoprecipitation, we find that BVRA does not bind or associate with ubiquitin (Fig. S2A). Additionally, we observed no changes in levels of Nrf2 in the presence or absence of BVRA both under basal conditions or in response to H_2_O_2_, sulforaphane or *t*-BHQ exposure (Fig. S2B). Next, we monitored Nrf2 levels in the nucleus of WT and BVRA^−/−^ cells upon induction of oxidative stress or with treatment with Nrf2 activators such as sulforaphane. While whole cell levels of Nrf2 are higher in WT cells with oxidative stress compared to cells lacking BVRA, the levels of Nrf2 in the nucleus are similar in both WT and BVRA^−/−^ MEFs under the conditions analyzed (Fig. S3C).

Lastly, we investigated whether binding of BVRA to Nrf2 is modulated by its substrate, biliverdin. We transfected HEK293 cells with GFP-Nrf2 and either GST-BVRA or GST vector, treated cells with biliverdin and conducted a pulldown assay. In the presence of biliverdin, interaction of BVRA with Nrf2 decreased as compared to cells which were not treated (Fig. 3*G*). Thus it appears that the presence of substrates for BVRA can influence its non-canonical functions.

### BVRA is a component of the transcriptional program of Nrf2-regulated promoters *in vivo*

BVRA has been reported to be a transcription factor regulating the expression of activating transcription factor 2 (ATF2) and HO-1 by binding to the AP-1 sites in their promoters (43, 44). Considering the physical and genetic interaction of Nrf2 and BVRA, we examined whether BVRA might potentially coregulate Nrf2 target genes. We performed a genome-wide ChIP-seq analysis of MEFs generated from previously described FLAG-BVRA transgenic mouse line (21) using FLAG and Nrf2 antibodies (Fig. 4*A-D*). The peak distribution demonstrated that approximately 10 % of the peaks were located in promoter regions with majority (approximately 40%) harbored in distal intergenic regions (10 to 50kb; Fig. 4*B*). Motif analysis of both FLAG-BVRA and Nrf2 ChIP-seq peaks identified bZIP motifs such as Bach1::Mafk motif and Nrf2 motif respectively (Fig. 4*B*). ChIP-seq peaks were annotated by nearest neighbor method to identify genes that both Nrf2 and BVRA potentially regulate. Approximately 43% of the genes in the annotated ChIP-seq data had both BVRA and Nrf2 peaks (*SI Data Table 2*). We further overlaid the BVRA-Nrf2 gene set with RNAseq-data from WT and BVRA^−/−^ cells. K-means clustering of the Nrf2-BVRA gene expression identified two clusters that are predominantly enriched for pathways that are dysregulated in neurodegenerative diseases (Fig. S3). Further, Gene Set Enrichment Analysis (GSEA) analysis of BVR-Nrf2 associated genes revealed inflammatory pathways, heme metabolism and redox signaling (Fig. 4*E*).

**Fig. 4.**
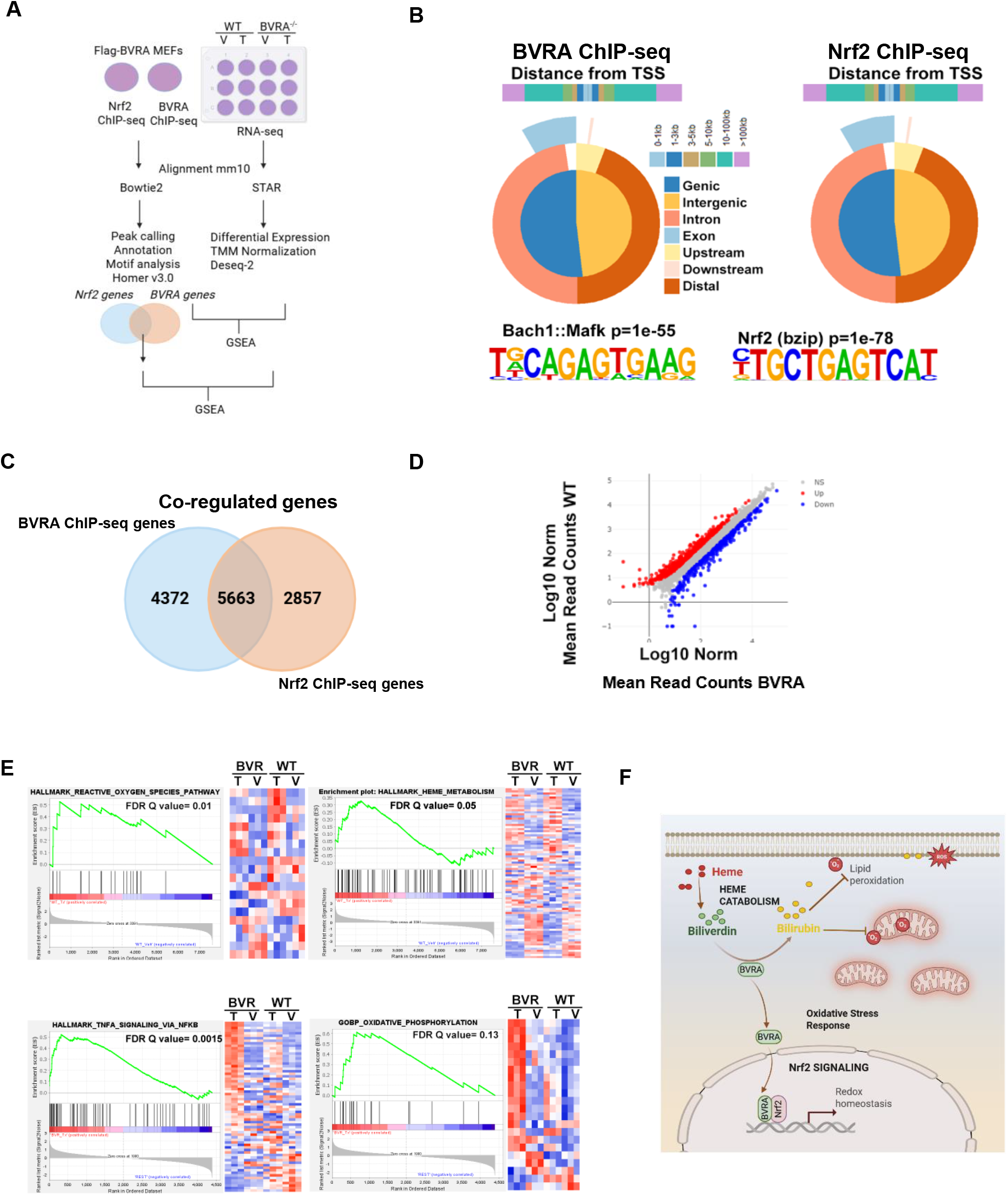
ChIP-Seq analysis reveals genes regulated by both BVRA and Nrf2. (*A*) Schematic representation of methods used for genomic analysis. (The data obtained from ChIP-seq was overlaid onto the RNA-seq data from the primary hippocampal neurons in vehicle treated (V) or H_2_O_2_-treated conditions (T) (*B*) Global representation of Flag-BVRA and Nrf2 peak distribution and Motifs. (*C*) Venn diagram showing overlap of genes associated with both BVRA and Nrf2 peaks. (*D*) MA plot showing differentially expressed readcounts between WT and BVRA^−/−^. (E) GSEA analysis of BVRA-Nrf2 associated genes from C. (*F*) Summary of canonical and non-canonical roles of BVRA. BVRA acts on heme-derived biliverdin to produce the antioxidant-cytoprotectant bilirubin, which scavenges superoxide generated from mitochondrial activity and other processes to maintain redox balance. In addition to this role in heme catabolism, BVR also functions as a serine-threonine and tyrosine kinase that modulates several signaling pathways. We show here that BVRA also regulates Nrf2 signaling, independent of its enzymatic roles.

Thus, interaction of BVRA with Nrf2 is physiologically relevant and fits in well with the observation that this association is increased in response to oxidative stress to increase the transcription of Nrf2 targets to maintain redox balance in cells

## Discussion

In this study, we show that BVRA, the biosynthetic enzyme for bilirubin, is an essential regulator of neuroprotective signaling by the master regulator of redox homeostasis, Nrf2. We showed previously that bilirubin is a potent antioxidant capable of scavenging superoxide (2). However, the role of BVRA in oxidative stress independent of its catalytic activity has not been explored (2). Using advanced genomic techniques, as well as evolutionary, mutational, genetic, and biochemical studies, we examined the non-canonical role of BVRA in the antioxidant stress response. sc-RNA-seq analysis of the brain revealed that a significant proportion of brain cell populations expressed BVRA but not HO-2, the precursor to BVRA in heme catabolism in the brain, indicating additional functions for BVRA distinct from its classical role in bilirubin production. Phylogenetic analysis revealed that although HO-2 and BVRA originated in the cyanobacteria in the context of phycobilisome synthesis in the light-harvesting complex, they were acquired at different points in time in the eukaryotes. Thus, they were co-opted and combined in a common heme catabolism pathway only in Metazoa. Using sequence conservation information from a multiple sequence alignment, we mutated key residues in BVRA and tested these mutants for BVRA activity. Further, the presence of a single exposed conserved cysteine within a CC motif at the extreme C-terminus in vertebrates suggests that this residue might be involved in the sensing of the redox status of the cell.

Amongst the mutants generated, the G17A mutant was severely compromised in the ability to produce bilirubin as this mutation affects its ability to bind the cofactor, NAD(P)H as reported earlier (21, 33). Interestingly, G17A BVRA rescued H_2_O_2_-mediated cell death, demonstrating a protective role of BVRA, distinct from its catalytic activity. Furthermore, unbiased RNA-seq analysis of hippocampal neurons derived from BVRA^−/−^ mice, identified Nrf2 signaling as a top downregulated pathway. In support of these findings, both WT and G17A BVRA bind Nrf2 and stimulate transcription of target genes. We also noted that in the presence of biliverdin, the substrate for production of bilirubin, the binding of BVRA to Nrf2 is diminished. This suggests a dual yet distinct role of BVRA with its additional ability to regulate redox metabolism via Nrf2. Although BVRA does not influence the nuclear translocation of Nrf2, it is essential for optimal function of Nrf2 as revealed by our RNA-seq and ChIP-seq data. This role of BVRA as a regulator that interacts with a transcription factor is closely paralleled by another catalytically inactive Rossman fold oxidoreductase harboring a from the same clade as BVRA, Gal80, which has been co-opted as a regulator that binds transcription factors (45).

Our studies revealed that multiple pathways are co-regulated by the BVRA-Nrf2-bound genes, which encompass a range of functions, including mitochondrial function, heme metabolism to cytokine signaling and oxidative stress response. Delineating this function of BVRA allows us to understand heme metabolism in the context of oxidant-induced aging and disease. While heme oxygenases, which catalyze the first step of heme degradation, are known targets for redox regulation, their overexpression leads to deleterious off-target effects including inflammation and cognitive decline (46) reducing its therapeutic efficacy (47). Likewise, while bilirubin itself acts as an endogenous and exogenous ROS suppressor, its effects are limited to distinct forms of ROS (2). While exogenous bilirubin may only be effective in curtailing excess O_2_^•−^, activators of BVRA expression may be able to regulate ROS in a more all-encompassing manner. The BVRA/Nrf2 interaction is relevant to a wide array of ROS-mediated pathologies. Suboptimal Nrf2 activity has been observed in neurodegenerative diseases such as PD and AD (48, 49). However, currently known activators of Nrf2 are predominantly electrophilic compounds that exert adverse side effects. In such a scenario, molecules that activate BVRA expression or stimulate its transcriptional activity are of significant therapeutic value. BVRA’s role in activating Nrf2 provides a new therapeutic signaling pathway that mitigates a broader array of oxidative species via multiple downstream effectors. Augmenting BVRA therefore promises to a more reliable and efficacious approach for diseases involving imbalanced Nrf2 activity. Additionally, modulating the Nrf2 pathway through BVRA may also mitigate ferroptosis, an iron-mediated form of cell death, which is observed in several neurodegenerative diseases as both BVRA and Nrf2 regulate iron disposition, through its metabolism or transport, and loss of either of these proteins result in accumulation of excess iron and vulnerability to oxidative stress (50–53).

Collectively, our findings reveal a previously unrecognized and crucial function of BVRA, which bridges two distinct and critical pathways, namely heme metabolism and Nrf2 signaling, to afford cytoprotective effects. Furthermore, BVRA orchestrates antioxidant stress responses in both the lipophilic and hydrophilic subcellular compartments of the cell through not only the production of bilirubin which directly scavenges O_2_^•−^radicals, but also through its role as a transcriptional regulator, respectively (Fig. 4*F*). Augmenting this latter pathway may offer therapeutic benefits in diseases characterized by redox imbalance. Both the enzymatic and non-enzymatic roles of BVRA participate in diverse aspects of normal physiology, and characterizing these activities and their cell-type-specific roles will likely reveal new paradigms for treating neurodegenerative diseases.

## Materials and Methods

### Plasmids/cDNA

Plasmids encoding GST-BVR (GST-BVRA) and GST-BVRB were generated by cloning cDNA encoding human BVRA and BVRB into pCMV-GST vector (Clontech/TaKaRa). Likewise, the myc-BVRA (myc-BVRA) plasmid was generated by cloning cDNA of human BLVRA into pCMV-myc vector. The GFP-Nrf2 plasmid was generated by cloning cDNA of Nrf2 into GFP-vector.

### Cell Culture

#### Mouse Embryonic Fibroblasts (MEFs)

WT, BVRA^−/−^, and Flag BVRA MEFs were isolated as previously described (2). In brief, embryonic day 14.5 (E14.5) embryos were obtained from timed BVRA^+/−^ matings. The pups were decapitated and eviscerated, after which the remaining portion was trypsinized and sheared. Isolated MEFs were plated in 2 wells of a 6-well plate and cultured overnight in DMEM (Gibco) supplemented with 10% FBS, 100 U/mL penicillin and streptomycin, and 2 mM glutamine at 37°C with 5% CO_2_. MEFs were then expanded to 6-well plates and transiently transfected with SV40T antigen (Addgene) and maintained until stably proliferative. DNA was isolated from the heads of the pups and genotyped to confirm the genotype of the corresponding MEF cell line. Data for WT and BVRA^−/−^ cells were drawn from experiments on at least two independent cell lines of each genotype.

#### Human Embryonic Kidney (HEK) 293 Cells

HEK293 cells were obtained from the American Type Culture Collection. HEK293 cells were cultured in Dulbecco’s Modified Eagle Medium (DMEM), 10% fetal bovine serum, penicillin/streptomycin (100 U/ml), and glutamine (2 mM) in an atmosphere of 5% CO_2_ at 37°C.

#### Primary neurons

Primary WT and BVRA^−/−^ neurons were isolated as previously described (2). In brief, hippocampi from embryonic day 18 (E18) embryos were isolated and washed with HBSS. Tissue was then incubated with 0.25% trypsin in HBSS for 18 min at 37°C in a conical tube, after which trypsin was inactivated with the addition of FBS. Tissue were then pelleted at 2000*g* for 5 min at 25°C, after which the supernatant trypsin/FBS was aspirated off. Tissue were then rinsed twice with HBSS, resuspended in HBSS, and then gently triturated through a fire-polished glass pipette to promote additional dissociation. Tissue was then filtered through a 70 μm nylon cell strainer to remove debris. The dissociated cells were pelleted at 2000*g* for 5 min at 25°C and then resuspended in neuronal culture media (composed of Neurobasal (Gibco) supplemented with 2% B-27 supplement and 2 mM glutamine). Cells were filtered once more through a 70 μm nylon cell strainer. Dissociated cells were then plated onto rinsed, poly-D-lysine-coated dishes. Four days later, the conditioned media was supplemented with fresh neuronal culture media. The day before dissections, plates were prepared and coated in 0.25 mg/mL poly-D-lysine overnight at 37°C. Just prior to plating, plates were rinsed three times with sterile water and once with neuronal culture media.

#### Transfection

HEK293 cells were transfected with Lipofectamine 3000 (ThermoFisher) as per the manufacturer’s instructions. Briefly, HEK293 cells were seeded in either 10 cm dishes (3×10^6^ cells) or in 60 mm dishes (1×10^6^ cells). The next day, transfection complexes were added dropwise to the plated cells. For cells in 10 cm dishes, 25 μg of DNA, 23.7 μL of Lipofectamine, and 50 μL of P3000 were brought to a volume of 1 mL in serum-free Optimem media (Thermo). For cells in 60 mm dishes, 5 ug of DNA, 8.25 μL Lipofectamine, and 10 μL P300 were brought to a volume of 500 μL in Optimem. Cells were harvested 48 hours after transfection.

Additional details of reagents and methods are available in the *Supplementary Section*.

## Acknowledgments

BDP, SHS and AAP were supported by the American Heart Association and Paul Allen Foundation Initiative in Brain Health and Cognitive Impairment (19PABH134580006). BDP and AAP by NIH/NIA 1R01AG071512. BDP also acknowledges support from NIH/NIA 1R21AG073684, NIH NIDA, grant P50 DA044123, funding from the Solve-ME foundation and the Catalyst Award from Johns Hopkins University. B.T. was supported by NIH AG077396, NS101967, NS133688, and the Department of Defense HT94252310443. M.C.O. and S.M.S. were supported by NIH P01CA236778. AAP was supported by The Valour Foundation, as the Rebecca E. Barchas, MD, Professor in Translational Psychiatry of Case Western Reserve University and the Morley-Mather Chair in Neuropsychiatry of University Hospitals of Cleveland Medical Center, and by the Wick Foundation, Department of Veterans Affairs Merit Award I01BX005976, NIH/NIA RO1AGs066707, NIH/NIA 1 U01 AG073323, the Louis Stokes VA Medical Center resources and facilities, the Lincoln Neurotherapeutics Research Fund, the Leonard Krieger Fund of the Cleveland Foundation, the Meisel & Pesses Family Foundation, and an anonymous donor. LMI and LA are supported by the funds of the Division of Intramural Research of the National Library of Medicine at the National Institutes of Health. We also acknowledge biorender.com, a service we used to design our schematic figures and graphical abstract, and Intelligenomica LLC for genomics support.

