## Supplemental File for "Biliverdin Reductase A is a major determinant of neuroprotective Nrf2 signaling"

### Supplementary Information
### Biliverdin Reductase A is a major determinant of neuroprotective Nrf2 signaling

Chirag Vasavda[a,1], Ruchita Kothari[a,1], Navneet Ammal Kaidery[b,c], Suwarna Chakraborty[d], Sunil Jamuna Tripathi[d], Ryan S. Dhindsa[e,f,g], Samaneh Saberi[h], Julia E. Lefler[h], Priyanka Kothari[i], Cristina Ricco[a], Kalyani Chaubey[j,k,l,m], Adele M. Snowman[a], Michael C. Ostrowski[h], Eugenio Barone[n], Lakshminarayan M. Iyer[o], L. Aravind[o], Sudarshana M. Sharma[h], Andrew A. Pieper[j,k,l,m,p,q], Bobby Thomas[b,c,r,s], Solomon H. Snyder[a,d,t,2], Bindu D. Paul[a,d,t,u,2]

*Solomon H. Snyder; *Bindu D. Paul



#### Supporting Information Text

#### Materials and Methods

##### Western Blotting

For western blot analysis, tissues were dounce homogenized at 4°C in lysis buffer (pH 7.4 solution of 50 mM Tris-HCl, 150 mM NaCl, 0.1% SDS, 0.5% sodium deoxycholate, and 1% Triton X-100) supplemented with protease inhibitors (Sigma). Lysates were then pulse sonicated and centrifuged at 16,000*g* for 15 min at 4°C. Fifteen micrograms of cleared lysate were run on a 4-12% polyacrylamide Bis-Tris gradient gel in running buffer (pH 7.3 solution of 50 mM MES, 50 mM Tris Base, 0.1% SDS, 1 mM EDTA) and then transferred to a PVDF membrane. Membranes were blocked with 5% milk in TBS-T (pH 7.6 solution of 16 mM Tris-HCl, 140 mM NaCl, 0.1% Tween-20) and incubated with primary antibodies in 3% bovine serum albumin (BSA) (w/v) overnight at 4°C. The following day, membranes were washed with TBS-T, and then incubated with secondary antibodies in 3% BSA for 1 h at 25°C. The following primary antibodies were used: rabbit anti-BVRA (Abcam ab180208; 1:1,000), rabbit anti-Nrf2 (Abcam ab62352; 1:1,000), rabbit anti-Nrf2 (Santa Cruz sc-722; 1:1,000), rabbit anti-GFP (Cell Signaling 2956; 1:1,000), mouse anti-GST (Abcam ab3416 1:20,000), mouse anti-myc (Abcam ab9106; 1:10,000), and mouse anti-β-actin (Santa Cruz sc-47778 HRP; 1:10,000). The following secondary antibodies were used: sheep anti-mouse IgG (GE Healthcare NA931; 1:10,000), and donkey anti-rabbit IgG (GE Healthcare NA934; 1:10,000).

##### Subcellular Fractionation

Cells were lysed in membrane prep buffer (10 mM Hepes pH 7.4, containing 1 mM EGTA, 1 mM dithiothreitol, 10% sucrose) supplemented with protease inhibitors (Sigma). After lysis cells were snap-frozen in liquid nitrogen and then thawed briefly in a cold-water bath, and then kept on ice thereafter. A 30G needle was then used to lyse cells 3-4 times. The cell homogenate was then filtered through a nylon mesh (pore size 75 μm) to remove any intact cells. 100μL of this sample was collected and set aside for the whole cell fraction. The remainder of the sample was centrifuged at 1,000 *g* for 10 minutes at 4°C. The supernatant was collected and set aside as the cytosolic fraction. The pellet was then re-homogenized in membrane buffer and spun down once again at 1,000 *g* for 10 minutes at 4°C. The pellet was then washed once more in membrane buffer by pipetting buffer over the pellet without homogenizing it. The cells were then centrifuged one last time at 1,000 g for 10 minutes at 4°C, and the supernatant was discarded. The pellet was re-homogenized as the nuclear fraction. The previously collected cytosolic fraction was then centrifuged at 11,000*g* for 20 minutes at 4°C and the pellet was discarded. All whole cell, cytosolic, and nuclear fractions were then pulse-sonicated to shear DNA and spun down at 16,000 *g* for 15 minutes at 4°C. The fractions were then processed for immunoblot analysis.

**BVRA enzymatic activity assays**

BVRA activity was assayed at pH 8.5 at 37°C. 250 nM of purified WT and mutant BVRA were separately incubated for 5 min at room temperature in a pH 8.5 solution of 50 mM Tris, NADP+, glucose-6-phosphate, and glucose-6-phosphate dehydrogenase (NADPH Regeneration System, Promega v9510). The samples were then placed at 37°C and spiked with varying concentrations of biliverdin IXα (250 nM to 20 uM). Reaction rates were determined by monitoring the change in absorbance at 442 nm over time. Enzyme activity was determined by the Michaelis and Menten (Michaelis Menten 1913). In brief, the absorbance of bilirubin was monitored over time for each concentration of biliverdin. The slope of each curve in the linear range was recorded and plotted against the corresponding substrate concentration. This data was then fit to the Michaelis-Menten equation to determine the $K_m$ and $V_{max}$ of the enzyme-substrate reaction (1).

**Methylthiazoletetrazolium (MTT) assays**

WT and BVRA$^{-/-}$ MEFs were plated at a density of 75,000 cells/well in a 24-well plate. After 24h, BVRA$^{-/-}$ cells were transfected with empty vector, WT-BVRA, and G17A-BVRA respectively. The following day, the culture media was supplemented with varying $H_2O_2$ concentrations (0 to 250uM) for 6 hours, after which the media was replaced with fresh media containing 0.5 mg/mL MTT for 1 h. The media was then removed before dissolving the reduced cellular MTT in DMSO. Cell viability was determined by measuring the absorbance at 570 nm and normalizing absorbances to the vehicle condition.

**Single-Cell RNA Sequencing**

Single-cell RNA-sequencing was performed as previously described (2). Briefly, whole brains from postnatal day 0 pups were dissected and dissociated for sequencing. Single cell RNA-seq libraries were constructed using the 10X Chromium Single Cell 3′ Reagent Kits v2 according to manufacturer’s descriptions and subsequently sequenced on a NovaSeq 6000. Reads were aligned to the mm10 genome using the 10X CellRanger pipeline with default parameters to generate the feature-barcode matrix. Downstream quality control and analyses on feature-barcode matrices were performed with Seurat v3. Genes that were not detected in at least 4 cells were excluded. Cells with fewer than 1,000 genes or more than 5,000 genes were also excluded from analysis. We also excluded cells in which greater than 15% of reads mapped to mitochondrial genes. The filtered matrices were log-normalized and scaled to 10,000 transcripts per cell. Using the variance-stabilizing transformation in the FindVariableFeatures function, we identified the top 2,000 most variable genes per sample. We then harmonized gene expression across datasets prior to clustering by identifying anchors between samples in each dataset using the FindIntegrationAnchors function and then computing an integrated expression matrix from these anchors as input to the IntegrateData function. We then linearly regressed the number of UMIs per cell and percentage of mitochondrial reads using the ScaleData function and performed dimensionality reduction using PCA. For each dataset, we selected the top 30 dimensions to compute a cellular distance matrix, which was used to generate a K-nearest neighbor graph. The KNN was used as input to the Louvain Clustering algorithm implemented in the FindClusters function. We chose a resolution parameter of 0.8 for clustering via Louvain. To annotate and merge clusters, we performed differential gene expression analysis on the integrated expression values between each cluster using the default parameters in the FindMarkers function

**RNA isolation, reverse transcriptase PCR (RT-PCR), and quantitative-PCR (q-PCR)**

RNA isolation and qPCR were performed as previously described (3). In brief, WT and BVRA$^{-/-}$ cells were treated with vehicle or 50 μM $H_2O_2$ for 8 h. Total cellular or tissue RNA was extracted using the RNeasy Plus Universal Kit (Qiagen) per the manufacturer's instructions. RT-PCR was performed with the SuperScript III One-Step RT-PCR System (Invitrogen), whereas q-PCR was performed with the TaqMan RNA-to-$C_t$ 1-Step Kit (Applied Biosystems).

**Immunoprecipitation**

HEK cells were co-transfected with the plasmids of interest as indicated above. Two days later, the culture media was supplemented with 10 μM MG-132 (Sigma M744) for 4 h. Cells were then lysed in immunoprecipitation buffer (1% Triton-X100, 50 mM Tris, 150 mM NaCl, 5% glycerol, 25 mM NaF, 1 mM $Na_3VO_4$) supplemented with protease inhibitors (Sigma) and placed on a shaker at 4°C for 15 min. The lysates were then centrifuged at 16,000*g* for 15 min at 4°C. IP samples were incubated with 1 mg of protein lysate and 40 μL of either Glutathione Sepharose 4B (GE Healthcare) or anti-FLAG M2 (Sigma) beads overnight at 4°C. 5% of each sample was set aside as input prior to addition of beads. The following day, beads were pelleted by centrifugation at 2,500g for 2 min at 4 °C. The supernatant was aspirated, and beads were washed 5 times in immunoprecipitation buffer by centrifugation, aspiration, and resuspension. Proteins were eluted from beads by boiling for 5 min at 100°C in LDS sample buffer (Thermo Fisher) and the eluates analyzed by Western blot as described above.

**Nrf2 ARE Luciferase Reporter Assay**

Nrf2 activity was monitored with a reporter cell line in which changes in Nrf2 activity are coupled to the expression of firefly luciferase (Nrf2/ARE Luciferase Reporter HEK293 Stable Cell Line). Nrf2 ARE HEK293 cells were infected with lentivirus encoding either empty vector, WT-BVRA, or G17A-BVRA were plated at 125,000 cells/well in a 24-well plate. The following day, cells were treated with 200 μM $H_2O_2$, 10 μM SFN, and 25 μM TBHQ for 6 h. Cells were then lysed in passive lysis buffer (Promega E1941) supplemented with protease inhibitors (Sigma) and let shake at 4°C for

15 minutes. The lysates were then centrifuged at 16,000 *g* for 15 minutes at 4°C and protein estimated using the BCA assay. 15 µL of lysate was mixed with 5 µL of luciferase assay substrate LARII (Promega) and the amount of light produced was measured using a luminometer for a 10 second integration time with a delay of 2 seconds. The luciferase measurement was normalized to protein and reported normalized to the vehicle condition.

**GST Firefly Dual Luciferase Assay**

WT and BVRA$^{-/-}$ MEFs were plated in a 24-well plate and transfected with NQO1 or GST ARE firefly luciferase and renilla luciferase. Two d post-transfection, cells were treated with $H_2O_2$, SFN, and tBHQ for 8 hours. Cells were then lysed in passive lysis buffer (Promega E1941) supplemented with protease inhibitors (Sigma) and let shake at 4°C for 15 minutes. The lysates were then spun down at 16,000*g* for 15 min at 4°C and protein estimated using the BCA assay. 15 µL of lysate was mixed with 5 µL of luciferase assay substrate LARII (Promega) and the amount of light produced was measured using a luminometer for a 10 second integration time with a delay of 2 seconds. Afterwards, 5 µL of Stop&Glo Reagent (Promega) was added and measured for 10 seconds with a delay of 2 s to estimate the amount of Renilla luciferase produced. The Renilla measurement and was used as the protein loading control to normalize the signal given by LARII. Data was normalized to the vehicle condition.

**Chromatin immunoprecipitation-sequencing (ChIP-seq)**

Chromatin immunoprecipitation (ChIP) assays were performed on cultured FLAG-BVRA mouse embryonic fibroblasts from 10 million cells per IP as described previously (4). The chromatin was immunoprecipitated with 10 µg anti-FLAG-M (Sigma Aldrich), 10 µg anti-Nrf2 (D1Z9C-XP Cell Signaling Rabbit mAB #1271) antibodies. 5 ng of the immunoprecipitated chromatin was used for ChIP-seq library preparation using NEBNext library kit (New England Biolabs) and sequenced in Illumina sequencer (50 million paired end reads per sample). Bowtie2 software (5) was used to map the sequence reads to the mm10 mouse genome. Peak calling and motif analysis were done with HOMER v3 software (6). Global annotation of peaks with respect to TSS and gene body was performed with the R Bioconductor package ChIPseeker (7).

**Computational analysis**

Homologous sequences of BVRA were retrieved using sequence-profile searches done with the PSI-BLAST program (8) against a curated database of completely sequenced genomes as well as a non-redundant protein database from the National Center for Biotechnology Information clustered down to 50%. Retrieved sequences were aligned with the Kalign2 (9) and MAFFT programs (10). For the MAFFT program alignment, the local-pair algorithm was combined with the following parameters: maxiterate, 3,000; op, 1.5; ep, 0.2. The aligned sequences were then used to perform phylogenetic maximum likelihood analyses using the FastTREE2 program (11). Cartoon structures were rendered with the Mol*(12) or PyMOL (The PyMOL Molecular Graphics System, Version 1.2r3pre, Schrödinger, LLC) software. Alphafold3(13) was used to model predicted interactions of BVRA with its NADPH, Biliverdin and Nrf2

**Statistical Analysis.** Results are presented as means ± SEM for at least three independent experiments. The sample sizes used were based on the magnitude of changes and consistency expected. Statistical significance was reported as appropriate. *P* values were calculated with Student's *t* test.

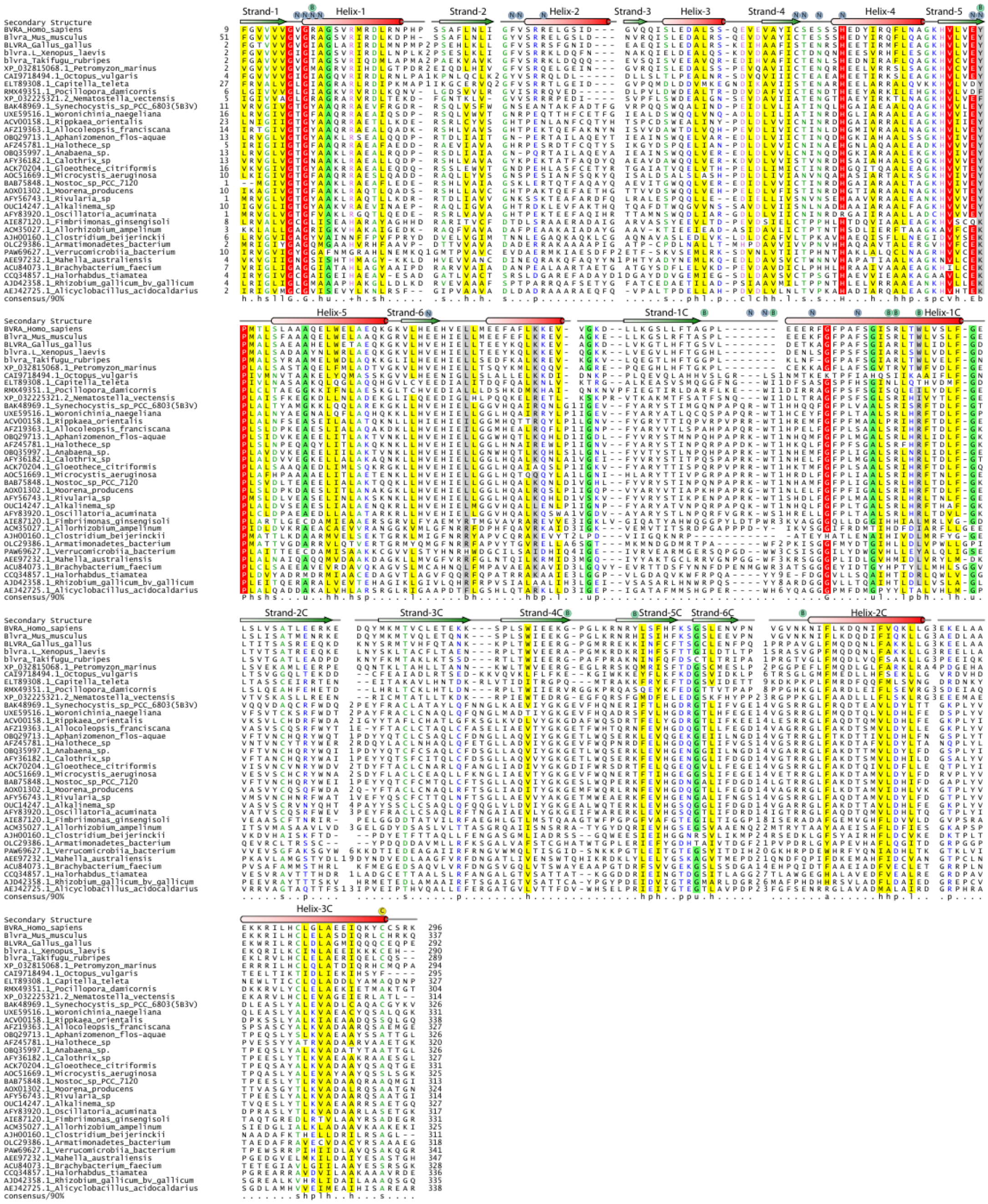

**Fig. S1. Domains in BVRA and alignment of BVRA from various species reveals conserved residues in the NAD(P)H and biliverdin binding sites.** Multiple sequence alignment of BVRA homologs from eukaryotes to bacteria. Proteins are labeled by their gene names or accession numbers and species names. Secondary structure is provided above the alignment with helices represented by cylinders and strands by arrows. Residue columns are colored by their consensus with absolutely conserved residues shaded red, h (hydrophobic), l (aliphatic), and a (aromatic) residues shaded in yellow; p (polar) residues shown in blue; c (charged) and + (basic) residues are shown in pink font; s (small) is shown in green font; u (tiny) is shown on a green background; b (big) is shaded gray. The NADPH binding residues (N) and biliverdin binding residues (B) were derived from the structure of the Synechocystis BvrA (pdb: 5B3V). The few additional residues that also bind NADPH in the Rat BVRA (PDB: 1LC0) are also included. C292 of Human BvrA is also indicated.

**A**

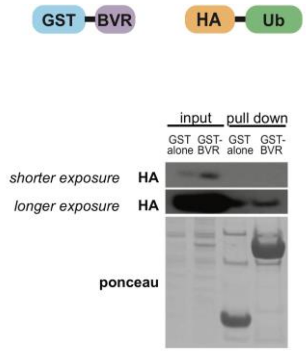


**B**

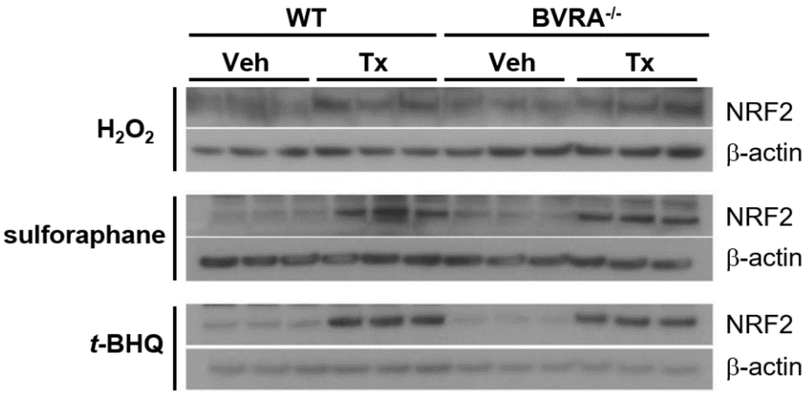


**C**

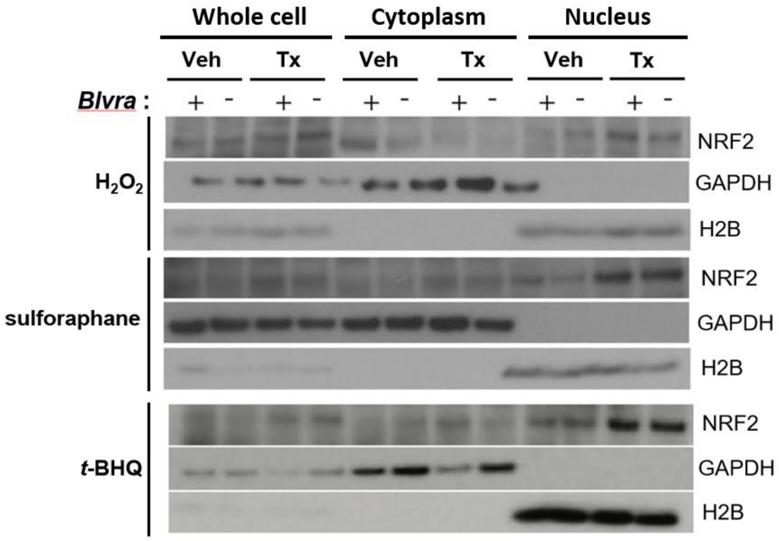


**Fig. S2. BVRA does not interact with ubiquitin or modulate either Nrf2 stability or its nuclear translocation.** (*A*) BVRA does not interact with Ubiquitin Provide details for each of these. (*B*) Ubiquitination of Nrf2 does not play a role in interaction with BVRA. (*B*) BVRA does not regulate the stability of Nrf2. (*C*) BVRA does not regulate the nuclear localization of Nrf2.

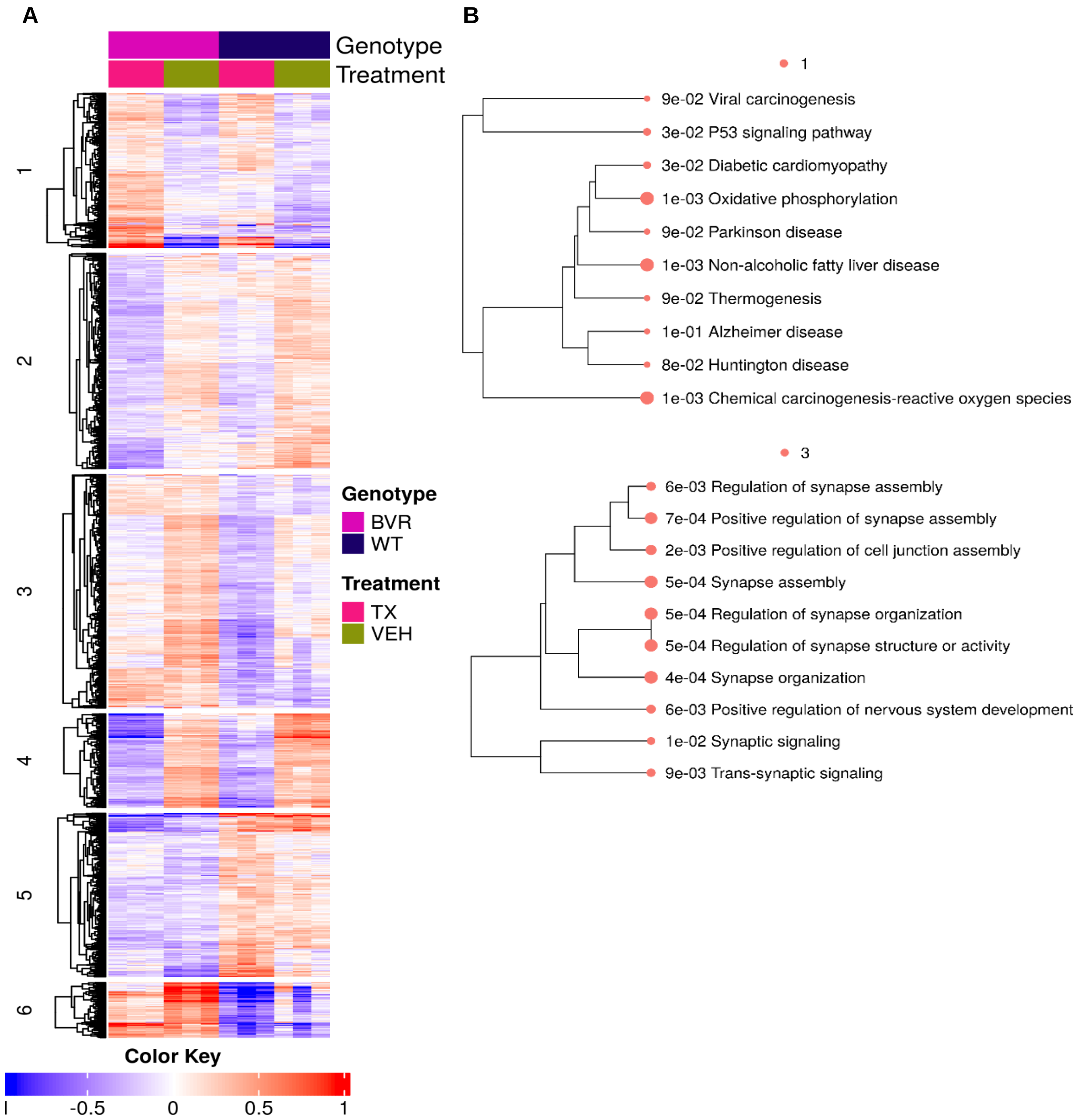


**Fig. S3. BVRA Nrf2 coregulated genes are critical in neurophysiology.** (*A*) K means clustering of expression of BVRA-Nrf2 associated genes in BVRA and WT (treated with either vehicle (PBS) or $H_2O_2$. (*B*) Pathway analysis of KEGG on cluster 1 and 3 that shows maximal enrichment with neuronal pathways.
